# Inference of ecological networks and possibilistic dynamics based on Boolean networks from observations and prior knowledge

**DOI:** 10.1101/2024.07.01.601264

**Authors:** Loïc Paulevé, Cédric Gaucherel

## Abstract

Being able to infer the interactions between a set of species from observations of the system is of paramount importance to obtain explaining and predictive models in ecology. We tackled this challenge by employing qualitative modeling frameworks and logic methods for the synthesis of mathematical models that can integrate both observations and expert knowledge on the system. After devising a formal link between ecological networks and the causal structure of Boolean networks, we applied a generic model synthesis engine to infer Boolean models that are able to reproduce the observed dynamics of a protist community. Our inference method supports optimization criteria to derive most parsimonious and most precise models. It is also able to integrate prior knowledge on the ecological network, adding constraints on impossible interactions, which is necessary to obtain realistic predictions. Such constraints may however prove to be too strict, in which case our method is able to conclude on the absence of a model compatible with both the observations and the input hypotheses. We demonstrated our methodology on experimental data of a protist system, and showed its ability to recover essential and sufficient ecological interactions to explain the observed dynamics.

## 1 Introduction

Understanding and predicting how are formed natural communities, often represented as species networks, is critical in ecology. Such communities have been studied by sequentially adding species or by invading an already formed community by a group of preselected species (Serván and Allesina, 2021). Such structural changes are highly conditioning the resulting interaction networks (Fukami, 2015). However, it is often highly difficult to guess the interaction network from field observations, except in rare lab experiments, thus suggesting developing inference methods aiming at identifying the interaction network leading to specific observed interactions (Milns et al., 2010; Pocock et al., 2021). Such networks are mandatory to anticipate the community dynamics (Veloz, 2019; Gaucherel et al., 2023).

Whereas many models of species community have been developed (Serván and Allesina, 2021; Baldan et al., 2018), inference methods are much less developed. Inference frameworks mainly build on Bayesian inference network aiming at identifying the interactions populating such networks Milns et al. (2010) or the interaction strength associated to a community (Pocock et al., 2021). They can even integrate expert knowledge and use supervised learning machines to improve the network inference (Auclair et al., 2017). Usually, such existing methods require a large amount of observations and knowledge on the studied system, in order to infer the interaction network and to calibrate the model. Does it concern the species (population) abundances and/or the interaction intensities (fluxes), statistical inference methods often use quantitative data for their parameterization.

To search for qualitative methods would be an original step in the way to identify ecological networks (Gaucherel et al., 2023; Dambacher et al., 2003). In such a case, the qualitative network of interactions (and interaction signs) would be inferred before going into the details of the interactions strength or probabilities, requiring more data. Indeed, several studies show that qualitative modeling frameworks, such as Boolean networks, greatly facilitate the inference of models of biological systems (Yu et al., 2004; Barroso-Bergadà et al., 2020), and can even enable reasoning over possible quantitative realization, without having to infer parameters (Paulevé et al., 2020; Gaucherel and Pommereau, 2019). In this paper, we demonstrated the feasibility of inferring ecological networks through the automatic design of Boolean dynamical models by integrating both observations of (community) system transitions and prior knowledge. To that aim, we first established connections between Boolean network (BN) theory and trophic networks, and show how global features such as the acyclicity and (in)sufficiency of trophic interactions can be translated into constraints over the architecture of a so-called influence graph of BNs (similar to the ecological interaction network). This enabled tailoring the generic method for Boolean network synthesis implemented in the software BoNesis (Chevalier et al., 2024), which relies on logic programming to automatically deduce Boolean models compatible with prior constraints on their architecture and possess specified dynamical properties, including reproducing given transitions, trajectories, and steady states.

We applied our methodology to the inference of a protist community network based on the observations gathered from one of the most exhaustive experiment ever in ecology (Weatherby et al., 1998) and has been well studied since then (Werner and Peacor, 2003; de Goër de Herve et al., 2022). Our inference strategy requires to first build a qualitative specification of dynamics, and specify constraints on the expected trophic networks (e.g., with a vertical hierarchy), related to expert knowledge. Then, employing BoNesis, one can deduce compatible Boolean networks and then the underlying trophic networks. We explored different optimization criteria, including identifying Boolean networks that reproduce dynamics as close as possible to the observations. In that sense, the inferred trophic networks are justified by the existence of a qualitative dynamical model that is able to replicate the observed transitions (Gaucherel and Pommereau, 2019).

In order to emphasize the flexibility of the framework to integrate prior expert knowledge on the ecological system, we performed the inference with (1) a zero-knowledge inference, only requiring acyclic trophic networks (2) prior on trophic networks from Weatherby et al. (1998), and (3) extended prior on trophic network from de Goër de Herve et al. (2022). This resulted in inferred community networks from increasing fits with the observed state transitions and the ecological knowledge. Models are required to have a perfect fit on observations (all system transitions are retrieved), and have minimal deviations, i.e., additional non-observed transitions. We finally discuss such improved inferences.

## 2. Materials and Methods

### 2.1 The protist community

As a case study, we focused on the seminal protist experiment published by University of Sheffield (Weatherby et al., 1998; Law et al., 2000; Warren et al., 2003, 2006). The experiment consisted in registering the successive states and transitions of a protist community made up to six species, chosen to build a trophic network involving various strategies and sizes. Selected species were *Amoeba proteus* (noted A; ∼ 200*µm*), *Blepharisma japonicum* (B; ∼ 200*µm*), *Paramecium caudatum* (P; ∼ 150*µm*), *Euplotes patella* (E; ∼ 140*µm*), *Colpidium striatum* (C; ∼ 70*µm*) and *Tetrahymena pyriformis* (T; ∼ 35*µm*), from largest to smallest sizes, feeding on unlimited bacterial resource. These species have been growing in a controlled (laboratory) environment and can only survive or die, without invasion. Their trophic and possible competition interactions, represented in fig. 1a, were first based on expert knowledge about their feeding habits, previously tested, and then on confirmed transitions observed between pools of species Weatherby et al. (1998); Warren et al. (2006).

**Figure 1:**
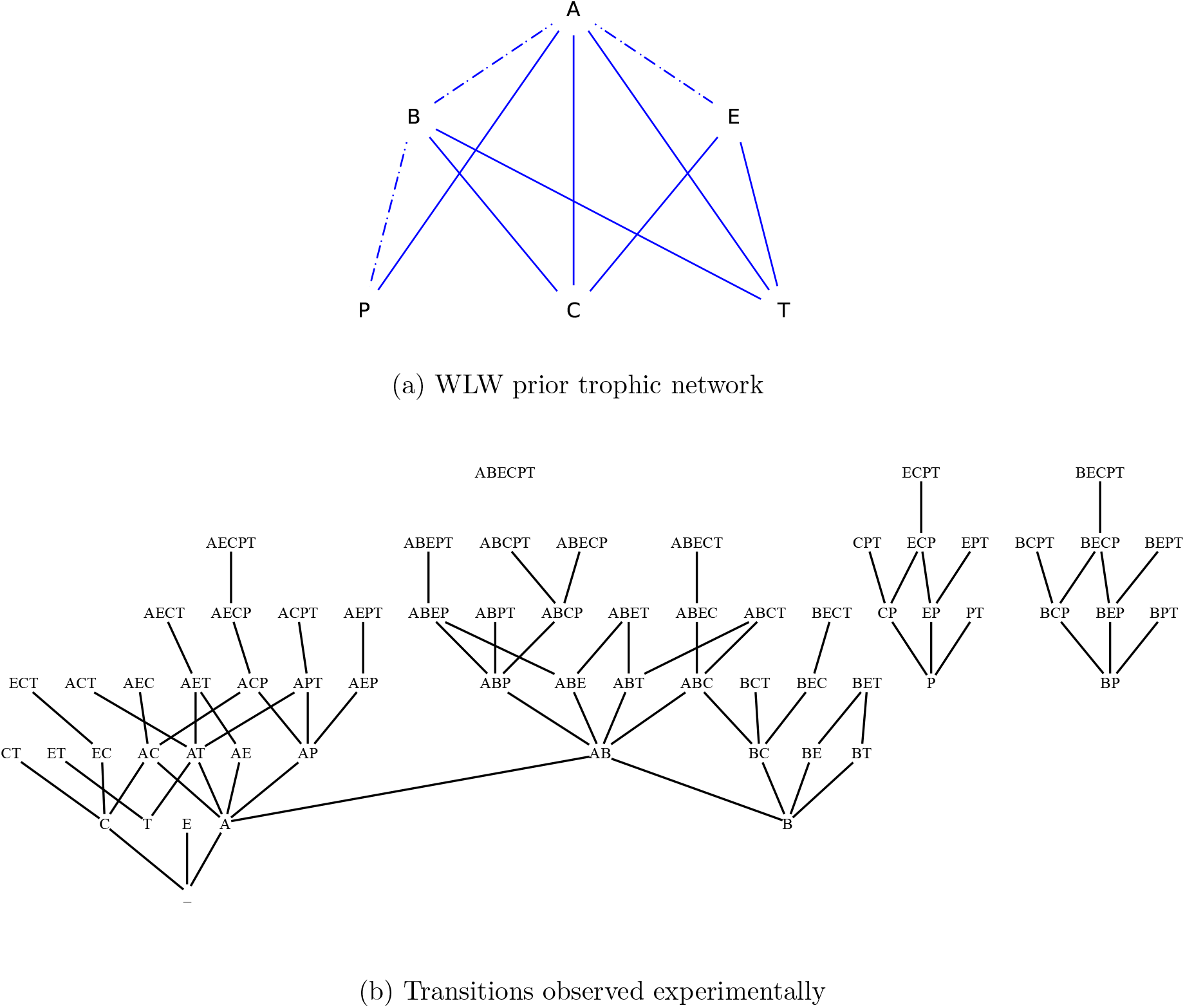
(a) **WLW prior trophic network** (Law et al., 2000) between six protist species feeding on unlimited bacteria resource (not represented). Dashed-dotted edges indicate insufficient trophic interactions. (b) **Qualitative transitions** derived from experimental observations. Chronology and reading direction is from top to bottom.

Each of the 63 combinations of protist species have been tested and replicated six times, with previous stabilized stock cultures and about a hundred individuals of each species. The resulting 378 microcosms were then sampled on day 17 and every 28 days until some chosen stopping criteria were reached. The observations stopped either when i) all species went extinct, or ii) only one species not able to persist alone on bacteria, or iii) only one species able to persist remained, or iv) species composition was unchanged for three successive samples. Most microcosms (93%) met one of these stopping criteria by 101 days in the original experiment (Weatherby et al., 1998). Here, we based our analysis on both the expert knowledge trophic network (Weatherby et al., 1998; Warren et al., 2003, 2006), as well as the full time series information available for reconstructing all qualitative time series and pathways of the protist community (Warren et al., 2003; de Goër de Herve et al., 2022). Indeed, we focused on presence/absence data of each species, previously confirmed by systematic scanning of the entire microcosm states after 20-30 times magnification, to draw a new trophic network with competition events in agreement with the observed transitions. The experiment revealed highly non-consistent community pathways, and no systematic final states or timing for transitions (Weatherby et al., 1998; Warren et al., 2003). Most of the time, the observed transitions were not reproducible, which appears quite usual, but is rigorously confirmed in this experiment. At least 32 out of the 63 experiments displayed multiple and different final compositions and pathways. Observed 71 transitions are summarized in fig. 1b. Each transition modifies the state of only one species. Furthermore, states with ingoing transitions but without outgoing observed transitions represent *steady states* of the system. They correspond to species B, P, T alone, BP, and total extinction (state). No observed transitions from state where all species are present: indeed, this state is not lasting, as many prey species are likely to rapidly disappear when all predator species are simultaneously present.

### 2.2 Boolean networks and influence graphs

*Boolean networks* (BNs) are discrete dynamical models which describe the change of activity state (on/off) of species using Boolean functions (Kauffman, 1969; Thomas, 1973). Formally, a BN *f* among *n* species is defined by *n* Boolean functions *f*_*i*_: 𝔹^*n*^ → 𝔹 for *i* = 1, …, *n*. Each function specifies the target Boolean state of the related species according to the Boolean state of the other species in the network. The state of all the species is represented by a *configuration*, which is a Boolean vector of dimension *n*, i.e., within 𝔹^*n*^.

The functions of a BN are typically represented using propositional logic (¬ is negation, ∧ conjunction, and ∨ disjunction). An example is given with its influence graph in fig. 2a.

**Figure 2:**
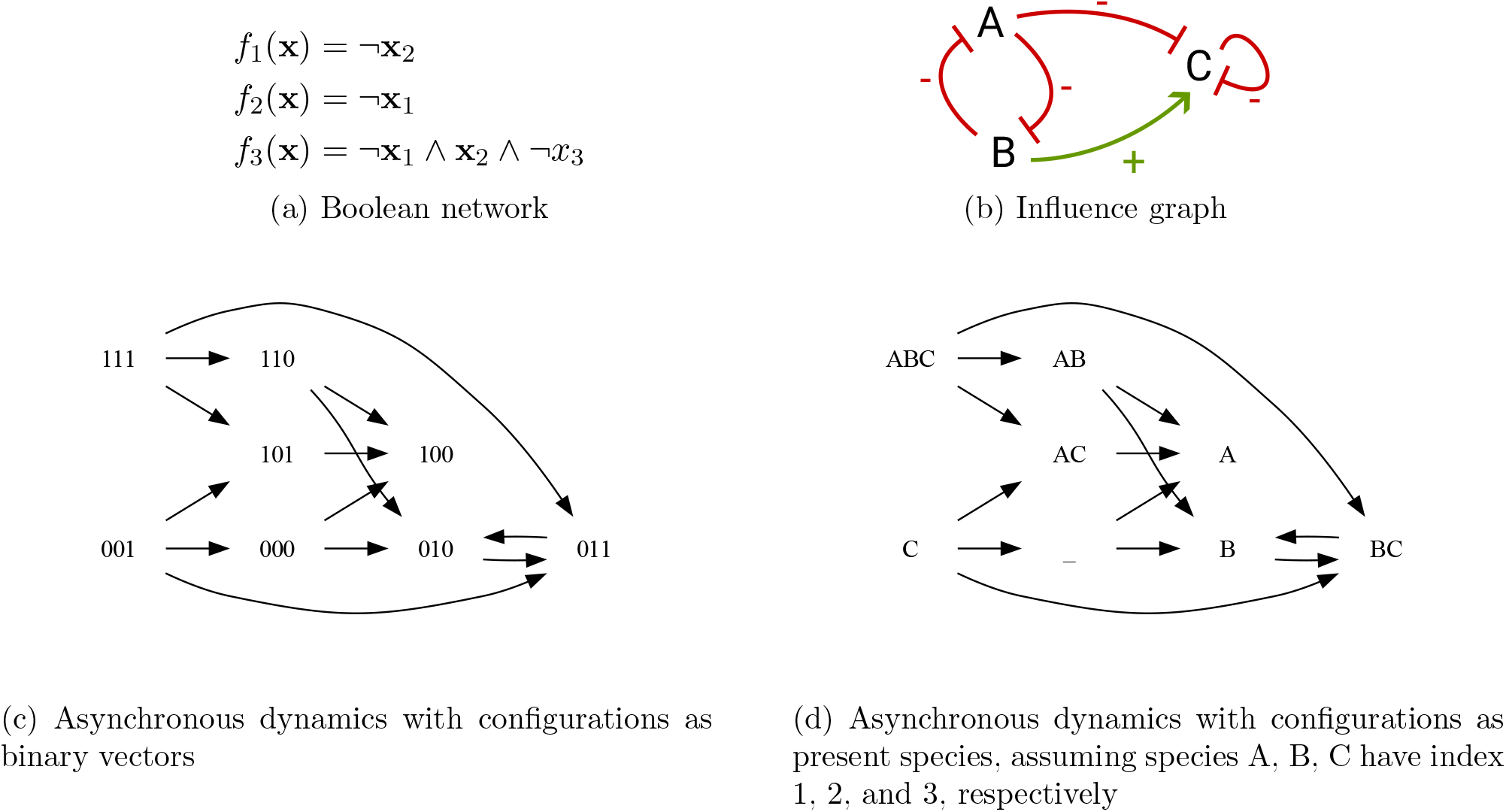
Example of BN with corresponding influence graph and asynchronus dynamics.

BNs are typically employed to predict successions of configurations and transitions, generating *trajectories*. Trajectories are computed from *f* by following an *update mode*. For instance, with the *synchronous mode*, all the species are updated in parallel, leading to a unique transition from any configuration **x** ∈ 𝔹^*n*^ to the configuration *f*_1_(**x**)*f*_2_(**x**) … *f*_*n*_(**x**). In the scope of this paper, we consider trajectories of BNs computed using the (fully) *asynchronous mode*, where only one species changes of state at a time. In this case, in general, several outgoing transitions are possible from a single configuration, leading to non-deterministic or undeterministic dynamics, sometimes called possibilistic (i.e., non-deterministic yet systematic). The non-determinism typically encompasses probabilities and stochasticity of transitions, as well as uncertainty on duration of state changes (Paulevé et al., 2020). BN dynamics can be captured by a *transition graph*, which is a directed graph DynA(*f*) between configurations 𝔹^*n*^ such that there is an edge from a configuration *x* to *y* if *y* is the result of the update of a single species *i* according to *f*_*i*_:

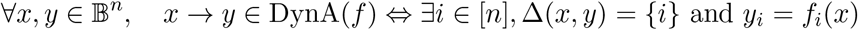

where Δ(*x, y*) = *{i ∈* [*n*] | *x*_*i*_ *x*_*i*_ ≠ *y*_*i*_} is the set of species whose state differ between configurations *x* and *y*.

Figure 2c gives an example of transition graph, with configurations represented as binary vectors. In this paper, we also represent configurations by the species that are at state 1, as shown by fig. 2d. An *attrator* is a non-empty set of configurations *A* ⊆ 𝔹^*n*^ that no trajectory can escape, and such that any pair of configurations are connected by a trajectory. Attractors can be composed of a single configuration, called *fixed points*, where the state of each species cannot change; or of multiple configurations, leading to sustained cyclic behaviors. They correspond to the terminal *strongly connected components* of the graph DynA(*f*) and model the long-term behaviors of the system.

The asynchronous dynamics of the BN of fig. 2a has two attractors: the fixed point{100}, and the cyclic attractor {010, 011}.

**Influence graph**. In general, the function *f*_*i*_ of a species *i* does not depend on the state of all the species of the network. The *influence graph* of *f* captures the (signed) dependencies between the functions of the species: there is positive (resp. negative) influence of *j* to *i* if there exists at least one configuration *x* in which the sole increase of *j* change the value of *f*_*i*_ from 0 to 1 (resp. 1 to 0):

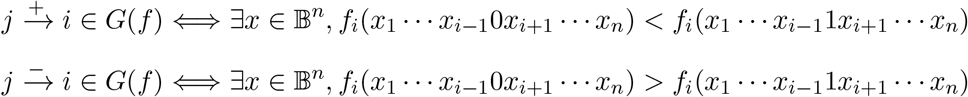

A BN *f* such that there exists *i, j* with both 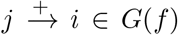 and 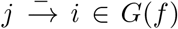 is *non-monotone*, otherwise it is *locally monotone*. In the scope of this paper, we do not consider non-monotone BNs and assume that for any species *j* it is either an activator 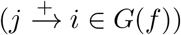, or an inhibitor 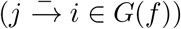 of another species *i*, or *j* has no direct influence on *i*.

An example of influence graph is given by fig. 2b. Positive edges are drawn in green, and negative edges in red with a bar arrow. The example BN is locally monotone.

**Note on figures**. In order to help distinguishing between the three type of graphs we are using, ecological (and trophic) networks are drawn with blue edges, influence graphs are drawn using green and red edges, and transition graphs are drawn with black edges. Note that in several figures, additional colors are used to highlight specific edges. All these graphs are directed, however, for the sake of readability, we omit arrow tips when direction is from top to bottom.

### 2.3 Inference of Boolean networks from structure and dynamics

We focus on the inverse problem of identifying BNs whose influence graph is included in a prior signed digraph, and whose asynchronous dynamics include the observed transitions and fixed points. More formally, the input of the inference problem is a signed digraph *𝒢*, a set of asynchronous transitions *T* ⊆ *{x → y* | *x, y ∈* 𝔹^*n*^, |Δ(*x, y*)| = 1}, and a set of configurations *S* ⊆ 𝔹^*n*^ which must be fixed points. The inference consists in finding BNs *f* such that *G*(*f*) ⊆ *G, T* ⊆ DynA(*f*), and for each *x ∈ S, f* (*x*) = *x*. Ecologists and modelers often use the Akaike or Bayesian information criteria for estimating the quality of a statistical model and its associated prediction error for a specific dataset (Burnham and Anderson, 2002). yet, such indices are not appropriate for our piece of work, in particular because they concern quantitative data and the absence of parameters in our qualitative models.

Many different BNs may fulfill these properties. We considered two measures:

- *sparsity*: we will focus on BNs requiring a minimum set of influences (interactions) between species. The minimalist criterion is computed with respect to set inclusions: a set of influences is minimal if there is no BN -using a strict subset of them while verifying the required dynamical properties. Note that there may exist several minimal BNs.
- *deviation*: we are interested in having models reproducing the observations as closely as possible. While we impose that a BN have *at least* the observed transitions, we can compute how many additional *unobserved* transitions are predicted by the model from the observed states. The less deviation, the better.

In this work, we employed the tool BoNesis (Chevalier et al., 2024, 2019), based on logic Answer-Set Programming (ASP) and clingo solver (Baral, 2003; Gebser et al., 2014). The tool is restricted to the inference of locally-monotone BNs, but can easily be extended to above-mentioned dynamical constraints and optimization criteria.

### 2.4 Methodology

Our general network inference strategy relies on the identification of BNs that match with observations and optional constraints on the structure of influences, and that are optimal with respect to sparsity and deviation. Then, from the identified BNs, we derive the underlying network of influence.

In this study, we required that compatible BNs verify *at least* the following dynamical properties:

- each observed transition (fig. 1b) is a transition possible in the asynchronous dynamics of the BN;
- the observed steady states correspond to fixed points of the BN: there must exist no outgoing transition from these configurations in the BN dynamics. The observed steady states are the five nodes in fig. 1b with in-going edges but without outgoing edges, i.e., B, BP, P, T, and no species ().

Besides the observations of the system, we investigated the impact of integrating expert knowledge on the structure of the influences. Specifically, in our case study, we considered *prior* trophic networks which delineate possible interactions between species: we imposed that any BN must have its underlying interaction network being a *sub-graph* of the prior network. The main effect of the prior knowledge is to forbid interactions between species: if there is a no edge from species A to B in the prior network, then, the Boolean function of B must not depend directly on A.

Here, we considered three inference scenarios:

1. without prior knowledge on trophic interactions, except that the trophic network must be acyclic, which we refer to as the *Neutral* model.
2. only using interactions referenced in the (early) prior trophic network of Law et al. (2000); (fig. 1a), which we refer to as the *WLW Prior*.
3. only using interactions referenced in the extended prior trophic network from de Goër de Herve et al. (2022), which we refer to as the *GeA Prior*.

Inferences have been run on a regular desktop computer (Intel Xeon 3.3Ghz with 64GB of RAM), and took between seconds to a couple of minutes depending on the cases.

The integration of prior knowledge on trophic networks for BN inference requires a formal characterization of their translation in terms of influence graphs of BNs. To our knowledge, this link has never been addressed so far.

#### From ecological networks to BN influence graphs

Trophic networks are directed graphs whose nodes are species and an edge from *i* to *j* means that *i* can be a predator of *j* and may feed on it. Moreover, the predation interaction may be qualified as *non-sufficient*, meaning that the prey alone is not sufficient to ensure the survival of the predator. For example, in the trophic network of fig. 1a, the species A cannot survive alone on species E, whereas it may survive alone on species T.

The BN defines for each species a function to determine its next state, either present (true), or absent (false). Essentially, if *i* is a predator for *j*, then we can deduce that:

- *f*_*i*_ can depend positively on *j*, i.e., there is positive influence from *j* to 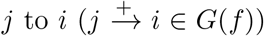 in its influence graph. This reflects that *i* may feed on *j*.
- *f*_*j*_ can depend negatively on *i*, i.e., there may be a negative influence from *i* to 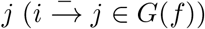 in its influence graph. This reflects that *j* can become absent because of *i*. More precisely, there may be a configuration where both *i* and *j* are present, which would then lead to *j* going extinct, whereas if *i* had not been there, it may survive.

In general, *f*_*i*_ can depend positively on itself, i.e., 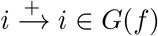, while having no self-trophic interaction. This is notably necessary if *i* cannot reappear in the community, i.e., in a state where a prey *j* is present, it may not be sufficient to go from absent to present; whereas in the same state, but where *i* is present, it will not disappear: *f*_*i*_(*i* = 0, *j* = 1) = 0 and *f*_*i*_(*i* = 1, *j* = 1) = 1; thus *f*_*i*_ depends positively on *i*.

These rules enable to translate a trophic network into a BN influence graph, which can then be used to specify the inference problem: find a BN using at most these influences and that is able to reproduce the observed transitions as close as possible. Figure 3 gives the translation of the WLW Prior as a BN influence graph.

**Figure 3:**
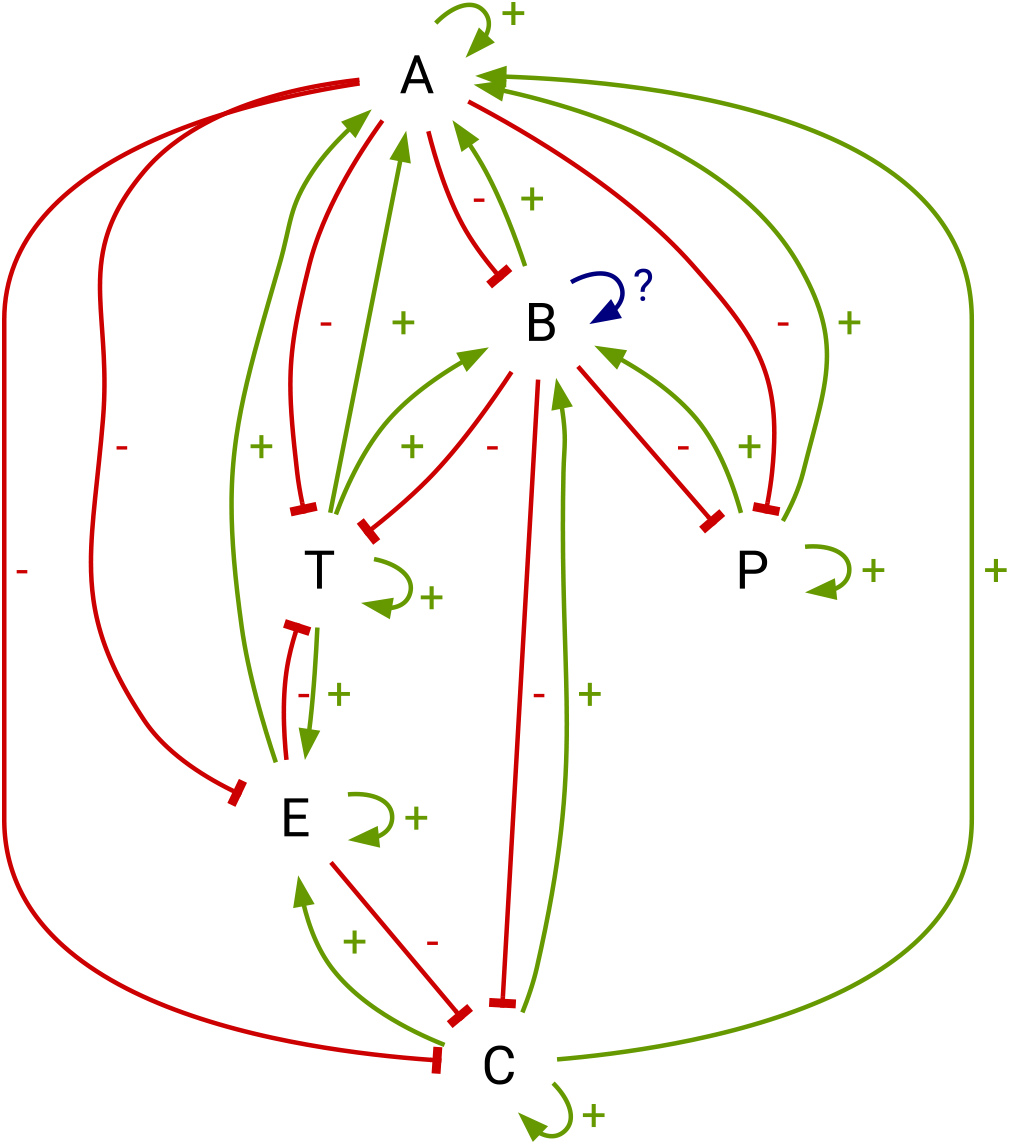
Influence graph corresponding to the WLW Prior (fig. 1a). Blue edges have an undetermined sign.

It must be stressed that the obtained prior influence graph delimits the set of candidate BNs: a candidate BN must employ *at most* the influences of the prior influence graph. In particular, it may employ fewer influences while being able to reproduce the observed dynamics.

The non-sufficiency constraints can also be integrated as constraints on the Boolean function themselves. Indeed, if the prior knowledge indicates that species A cannot survive alone on species B, we impose that the Boolean function of A cannot be true on configurations where all species but A and B are in state 0.

#### From BNs to ecological networks

The Boolean functions of the BN may involve both trophic and competitions between species. The influence graph of a BN captures the signed causal relationships between the species and their Boolean activation functions. From those signed digraph, we draw the corresponding ecological network as follows:

- there is an interaction from *i* to *j* in the ecological network if *f*_*i*_ depends positively on 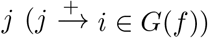 *i* feeds on *j*;
- there is an interaction from *j* to *i* in the ecological network if *f*_*i*_ depends negatively on 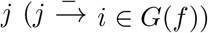: *j* is eaten by *i*.

Finally, an interaction from species *A* to *B* is marked as sufficient (plain line in figures 1 and 4) whenever it is sufficient that *A* and *B* are 1 in a state to satisfy the Boolean function associate to *A*, i.e., *f*_*A*_(*A* = 1, *B* = 1) = 1. If the condition is not sufficient, we mark the interaction as insufficient (dashed line).

**Figure 4:**
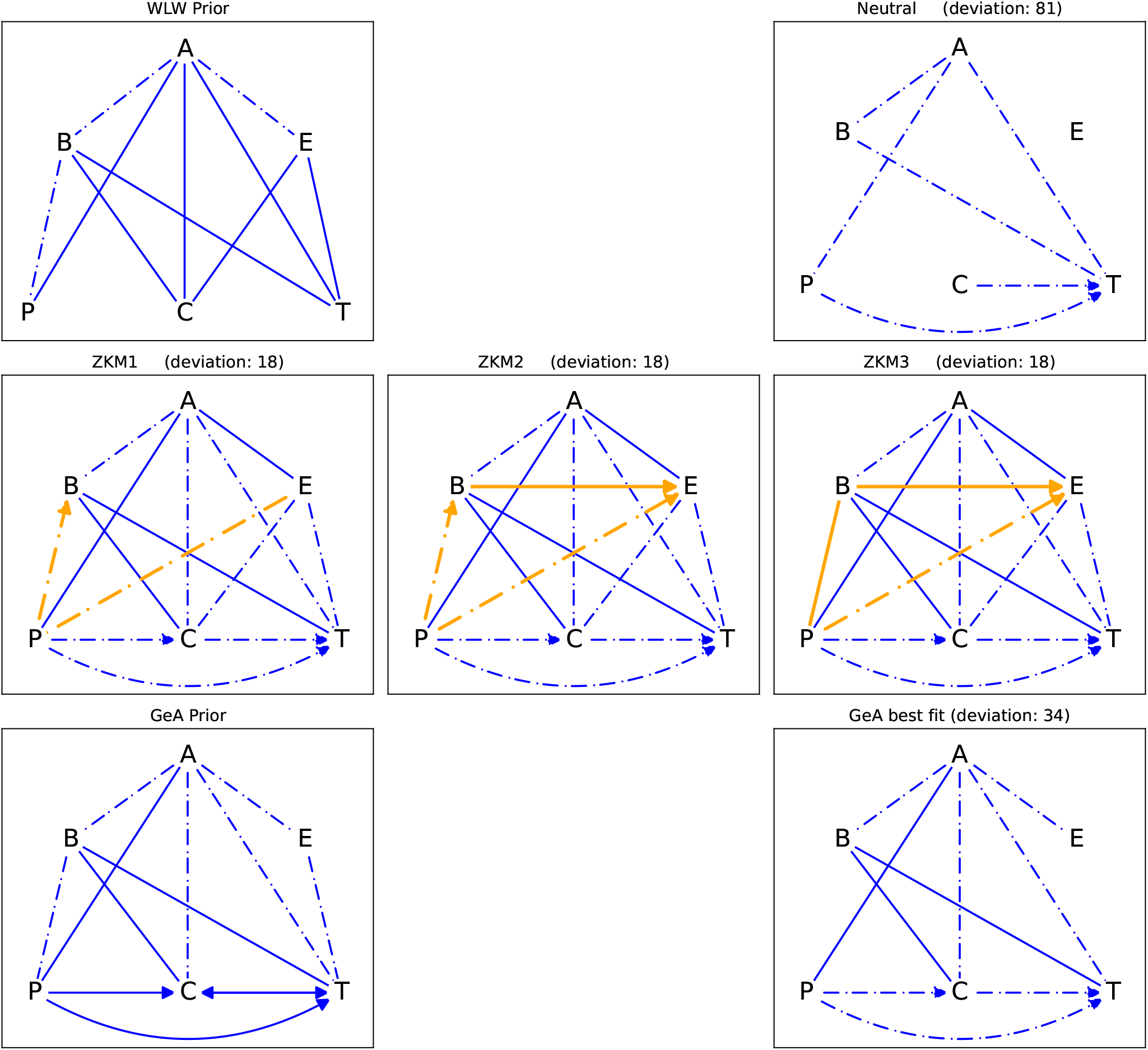
Prior and inferred ecological interaction networks. Unless specified, edge direction is towards the lower; there are no edges with default direction overlapped by a directed edge. Dashed-dotted edges represent predation that are not sufficient for the predator to survive only with this prey (starvation)… Bacteria are not displayed.

**Figure 5:**
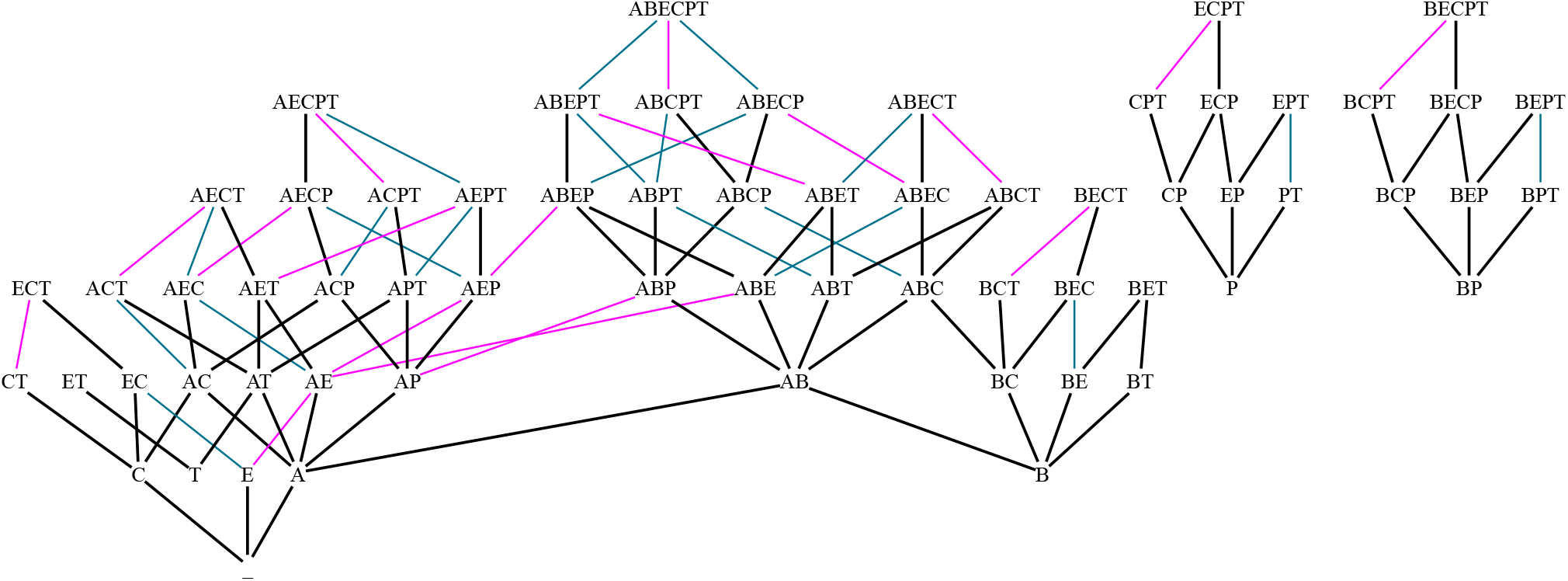
Transition graphs of inferred models. Transitions in black plain lines are the observations and are shared by all the models; in magenta are the 18 additional transitions of the ZKM3 and GoA models; in (dashed) teal are 16 additional transitions of the GoA model. Transitions of the Neutral, which include GoA transitions, are provided fig. A.

## 3 Results

We present the results of the inference of BNs from the observed transitions of the protist system and the different levels of prior knowledge on the trophic network between the six species. Recall that all the inferred BNs are able to reproduce the observed transitions and steady states.

### 3.1 Inference of Boolean models of protist systems from zero knowledge

We inferred BNs that can reproduce at least the observed transitions of the protist system, without *a priori* on the ecological network, except that this latter must be acyclic. Therefore, any possible interaction is considered, as soon as no species is directly or indirectly a prey of itself.

We inferred the sparsest (neutral) model, which in our case was unique. It shows the minimal interactions that are needed to be able to reproduce (at least) the observations. It depicts trophic interactions already close to the WLW prior trophic network, with the notable addition of the predictions of T by B, C, and P. The transition graph of the Neutral model contains all the observed transitions with the same observed steady states, alongside with 81 additional transitions (deviation). *ZKM1/ZKM2/ZKM3* are alternative inferred models that minimize deviation while reproducing the observations and having an acyclic ecological network. Their dynamics include the observed transitions and 18 unobserved transitions. It is the minimum achievable with acyclic ecological interactions, and observed transitions and steady states. The ZKM models differ only by the interactions between B, P, and E. Notably, ZKM1 and ZKM2 involve the predation of B by P, while ZKM3 needs the reverse. Compared to WLW Prior and Neutral, ZKM3 (and ZKM2) involves the predation of E by B and P. Considering each species size and their ecology, ZKM3 is the most realistic, as all interactions are compatible with a decreasing size from predators to preys.

Hence, even without *a priori* on the trophic interactions, the transitions and steady states are sufficient to identify plausible and sparse trophic networks by the means of Boolean network inference.

### 3.2 Inference with expert knowledge from WLW Prior

The second inference was performed by constraining the ecological network to be contained within the WLW Prior (fig. 4), including indicated non-sufficiency of interactions. We ignored constraints related to the sufficiency of trophic interactions to maintain a species, in order to consider more candidate models.

It resulted that *no model* using exclusively those pre-determined ecological interactions can reproduce all the observed transitions and steady states. The reason lies on the fact that in the trophic network, T is not related to C, and P. However, from the observations, T can survive alone in steady state, but in presence of C or P, T goes extinct. Indeed, the following transitions are given (see also fig. 1b):

- *CT → C*
- *CPT → CP*
- *PT → P*

It is therefore impossible to explain these above transitions without introducing additional interactions, thus horizontal predations or competitions. Note that this was indeed captured by the Neutral and ZKM models.

### 3.3 Extended WLW Prior

De Goër de Herve et al. (2022) proposed an extended trophic networks, coupled with potential competition, reproduced as “GeA Prior” in fig. 4. Using this extended ecological network, we identified automatically one (unique) model, minimizing its deviation from observations. This model, whose trophic network is shown in fig. 4 as “GoA best fit”, introduces 34 unobserved transitions. It is the best fit model while using only the interactions listed in the GeA Prior. Its deviation is larger than ZKM models. Compared to ZKM3, the model misses numerous interactions involving E, and the predation of P by B. Moreover, the predation of E is sufficient for A to sustain in ZKM models, while it is declared as insufficient in the GoA best fit model.

This model shows a trophic network rather close to the WLW prior, at the exception to the species T, that is here predated by P and C, which enables to reproduce the observed transitions related to the extinction of T in their presence. Hence, again, the absence of such trophic interactions made impossible the inference of a compatible model using only WLW Prior.

## 4 Discussion

### 4.1 Case study

The WLW Prior, and Neutral and GeA model present interactions more or less vertical, in terms of ecology and body sizes, leaving more or less room to the non-trophic (e.g., competition and other horizontal) interactions. Also, some models infer without *a priori* on the ecological network appear unrealistic (ZKM1 and ZKM2) as they infer “reverse predations” (smaller preys possibly eating larger predators). It is well know in aquatic ecology that the mouth size in constraining the prey size, thus of increasing values, but the reverse situation might happen in case prey adults sometimes feed on predator juveniles (Yasuyuki Choh and Janssen, 2012). Here, we often observed reverse role of E species, which is not so surprising considering only its comparable body size to P species (Weatherby et al., 1998; Warren et al., 2006). For this reason, this might be experimentally verified. We also observed here that T and E species often appear to be discriminant (ZKMs) and should be studied.

Our explorations also confirm that one single interaction may drastically change the computed dynamics, yet not necessarily the deviation to observations, as in the ZKM models. Similarly, we note here the importance of what could be called the “interaction status”, namely the ability to sustain the predator species (plain edges) or not (dotted edges), and the fact that some preys may be preferred or not (among several) by a predator species is crucial (e.g., ZKM2, species A). It might be carefully verified experimentally, as it may change the fate of the community by systematically removing a species before another one (and so forth) (Fukami, 2015; de Goër de Herve et al., 2022). For this reason too, the model deviation itself should not be regarded as the single discriminant criterion of inference, as the expert knowledge (on species behavior) may sharply change the inferred network too. And one (carefully selected) transition would be sufficient in refuting an inferred model, which is reassuring.

All the model we infer are able, by design, to mimic all the observed transitions, including the steady states. The minimum number of additional transitions, we qualified as deviation, depends on the *a priori* on the network structure One can observe in section 3 (and fig. A) that most of the additional transitions form so-called diamonds of transitions. E.g. starting from BEPT, the observations only relate transitions from BEPT to BEP to BP, and BPT to BP. The ZKM3 model adds the transition from BEPT to BPT. In qualitative dynamics, these diamonds typical reflect *concurrent* events: from BEPT, species E and T eventually disappear. If these events are independent of each other, they can disappear in any order, generating diamond-shaped transitions. As the asynchronous dynamics of BNs do not capture features related to speed or priority, one can indeed expect that there is not enough information to enforce a specific ordering in species disappearance. The only possibility to have more controlled transitions, i.e., having more complex causal structure to explain that a transition, is possible but not another from the same state. Thus, the importance of the prior ecological network, which, by denying direct interactions (those not referenced in the prior network), prevent deriving aberrant causal relations to fit as much as possible with the data.

### 4.2 Inference methodology

We demonstrated here an interactive method for generating hypotheses on underlying mechanisms driving ecological systems by the means of inference of qualitative models. To automatically infer (without or with little expert knowledge) ecological interaction networks becomes today mandatory in many observed ecosystems (Pocock et al., 2021; Milns et al., 2010). Our qualitative and possibilistic method (i.e., asynchronously exploring all possible transitions) appears here capable of identifying several interaction networks responsible for an observed dynamics, among which some are minimalist and display low deviations between dynamical observations and computations. In this study, we showed a complementary and parsimonious approach, requiring no quantitative data and little expert knowledge on the studied system (e.g., (Gaucherel et al., 2023; Gaucherel and Pommereau, 2019; Paulevé et al., 2020)). This enables to efficiently test several interaction networks fulfilling the observed dynamics, and derive candidate models that can then serve as a basis to derive more fine-grained and quantitative models.

Without specific prior knowledge on the trophic network, our approach automatically suggests the simplest explanation for the observations, i.e., using the least trophic interactions or least transitions as possible. This can constitute a first working hypothesis on the underlying ecological model, without the need of quantification. The inferred trophic interaction may then be validated or refuted, possibly by additional experiments and observations. This new expert knowledge can then be integrated, and a new inference process will return new models satisfying these additional constraints, in an iterative procedure (Baldan et al., 2018; Auclair et al., 2017). Our inference framework can also conclude to the absence of models satisfying both desired dynamical and structural properties. This typically indicates that the prior knowledge or the qualitative interpretation of data must be revised to obtain a comprehensive logical explanation of the systems dynamics.

We believe that community ecology largely requires such inference model for identifying the species networks at play in the field (Serván and Allesina, 2021; Fukami, 2015). This discipline is on the way to develop powerful inference, often based on frequentist and Bayesian models (Milns et al., 2010; Pocock et al., 2021). Yet, they often require a large amount of data and expert fine tuning rarely available in ecology and environmental sciences. A promising perspective would be to combine purely qualitative approaches to delineate a first ensemble of candidate models based on logical reasoning, before employing more fine grained modeling approaches to incorporate more information related to temporized time and species abundances.

## Acknowledgments

We warmly thank Mathieu de Goër de Herve who developed the first model from the discrete-event family (de Goër de Herve et al., 2022). We also thank Franck Pommereau, Colin Thomas and Maximilian Cosme for fruitful discussions on early versions of this work. LP was supported by the French Agence Nationale pour la Recherche (ANR) in the scope of the projects “BNeDiction” (grant number ANR-20-CE45-0001) and “REBON” (grant number ANR-23-CE45-0008).

## Author Contributions

LP and CG conceived the ideas and designed methodology; LP performed computational experimentation and CG performed analysis; LP and CG wrote the manuscript.

## Data and code availability

Data and code for reproducing the inferences are available at https://github.com/pauleve/bonesisprotists. The inference has been performed using BoNesis v0.6.5, available at https://bnediction.github.io/bonesis and archived at https://doi.org/10.5281/zenodo.10891199.

## A. Supplementary Materials

**Figure A:**
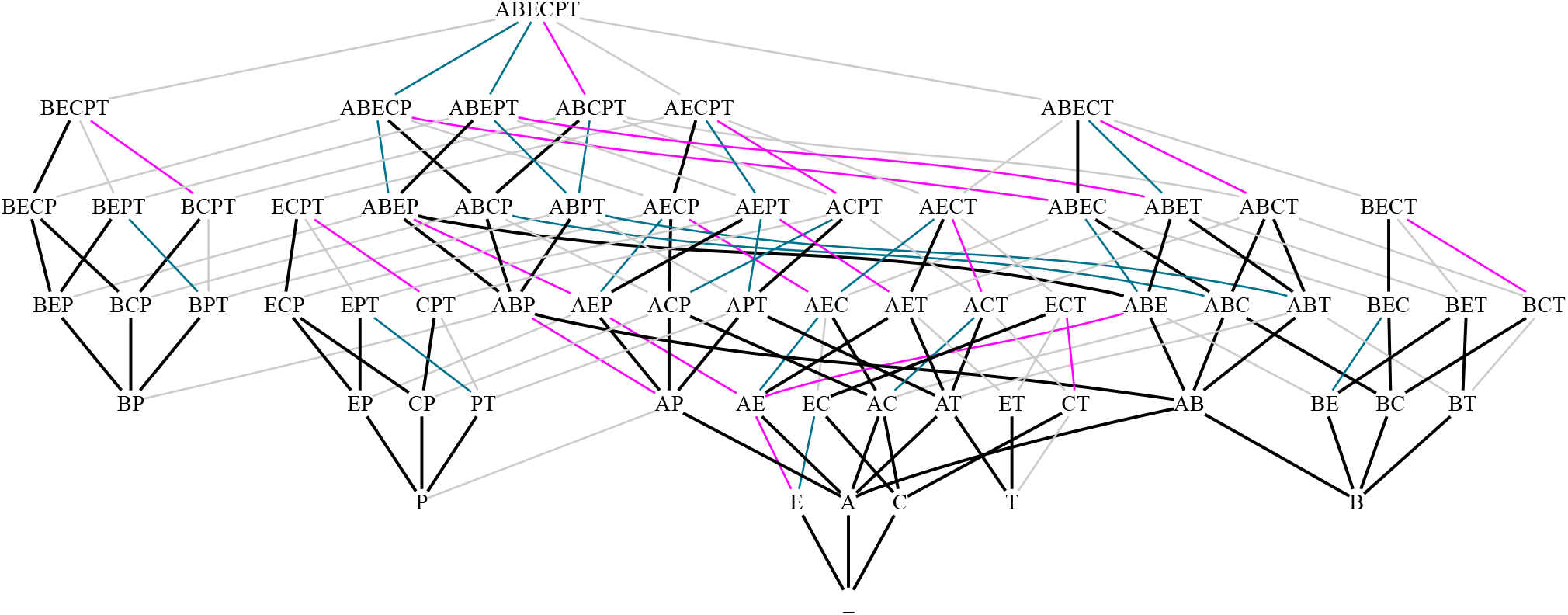
Transition graphs of inferred models, including Neutral. Transitions in black plain lines are the observations and are shared by all the models; in magenta are the 18 additional transitions of the ZKM3, GoA, and Neutral models; in (dashed) teal are 16 additional transitions of the GoA and Neutral model. In gray are the 47 additional transition of the Neutral model.

### A.1 Inferred Boolean networks

Neutral model:

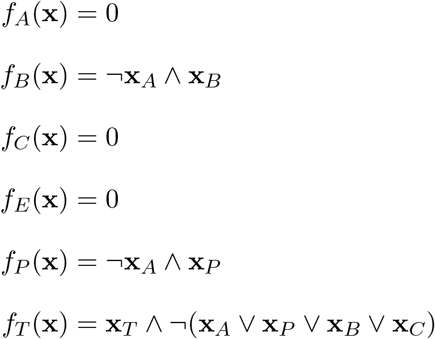

ZKM1:

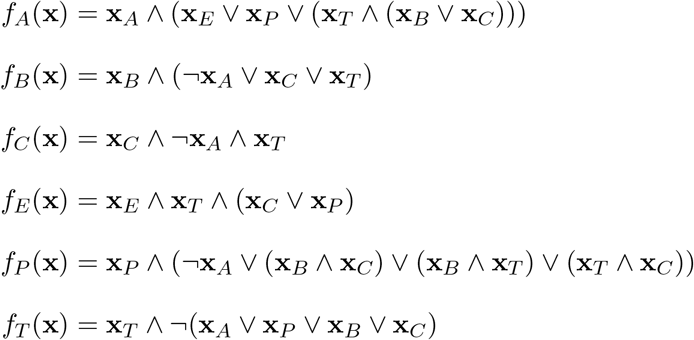

ZKM2:

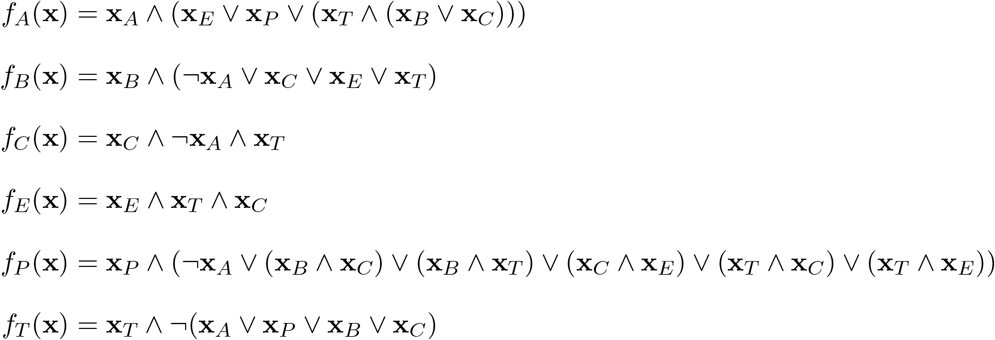

ZKM3:

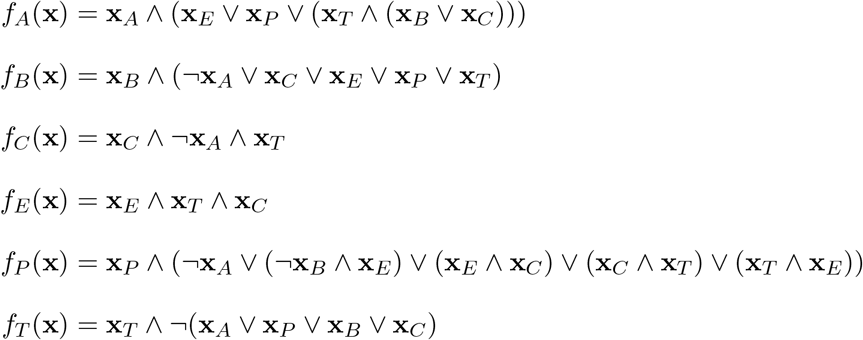

GoA best fit:

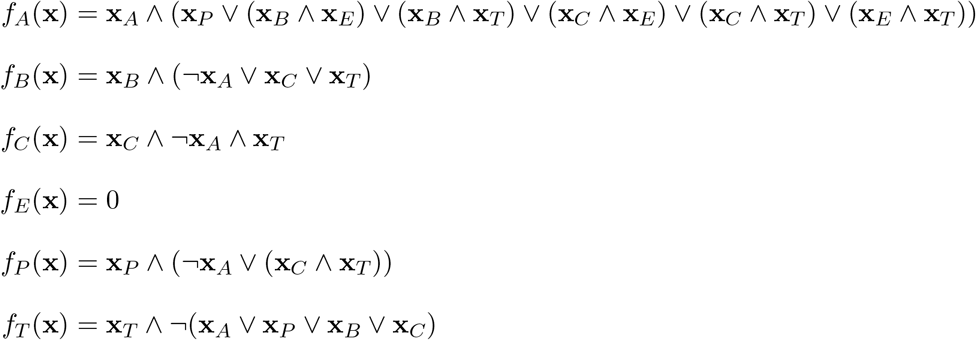

